# Horizontal transfers can reveal ghost birth events and complete species trees

**DOI:** 10.64898/2026.10.12.758438

**Authors:** Enzo Marsot, Bastien Boussau, Damien M. de Vienne

**Author notes:** Corresponding author: Enzo Marsot.

## Abstract

Extinct and unsampled species vastly outnumber the species typically included in phylogenetic studies. Most of these so-called “ghost lineages” are unknown and, among them, those that are extinct may have left no trace in the fossil record. Horizontal gene transfers (HGTs), however, provide a potential source of information about these missing lineages, because transfers can connect sampled ancestral species to lineages outside the sampled species tree. Such transfers may leave observable signatures in gene trees. Here, we formalize this signal under idealized models of duplication, transfer, and loss, and characterize the information that can be retrieved about the complete species tree (containing ghost and non-ghost lineages) thanks to sampled gene trees and sampled species trees (containing only the non-ghost lineages).

We establish four identifiability results, assuming an infinite number of gene families and errorless dated gene trees. First, transfers identify the lineage through time (LTT) curve of the complete tree, averaged over each time interval induced by the nodes of the sampled species tree. Second, in the presence of transfers, the distribution of node heights in sampled gene trees uniquely determines the internal speciation times of the complete species tree, including ghost speciations. Third, these hidden speciation times suffice to restore identifiability of birth-death-sampling parameters, through a reduction to a recent theorem of Dieselhorst and Stadler. Finally, when duplication, transfer, and loss rates are known constants and the transfer rate is non-zero, the distribution of sampled gene trees almost surely uniquely determines the topology and branch lengths of the complete species tree, including all ghost branches.

These results provide a theoretical basis for using signatures left by HGTs in gene trees to recover information about extinct and unsampled lineages, and suggest ways in which this signal could be exploited in practice.

## Introduction

Ghost species are species that are either extinct without having left traces in the fossil record, or extant but undiscovered or simply unsampled, *i*.*e*., not included in a given analysis [29, 28]. Due to the speciation and extinction events that have continually occurred since the beginning of life on Earth, we expect that extinct species vastly outnumber extant species [18]. Additionally, most clades are thought to be incompletely described [16, 13, 4, 8, 11, 12, 3], and even described species are often excluded from analyses for technical reasons or because they fall outside the scope of a study. Consequently, ghost species represent a major fraction of the diversity relevant to phylogenetic analyses.

In molecular phylogenetics, species trees are typically reconstructed from extant, sampled species and therefore contain information only about the ancestry of the sampled lineages. Assuming that all extant species are sampled and that the tree is generated under a constant-rate birth-death (CRBD) process with speciation and extinction rates *b* and *d*, these reconstructed trees allow the birth-death parameters to be identified. This property is lost when extant species are incompletely sampled: if the leaves constitute an unknown, uniformly sampled fraction *ρ* of all extant species, the resulting constant-rate birth-death-sampling (CRBDS) model is unidentifiable from reconstructed phylogenies: distinct triplets of speciation, extinction, and sampling parameters (*b, d, ρ*) may induce the same distribution of reconstructed trees and hence cannot be distinguished from these data alone [20, 22, 17, 14]. Additional information can restore identifiability. Fossils, for example, provide such information through the fossilized birth-death framework [21, 9], and can in some settings resolve this non-identifiability [31]. However, many clades, especially bacteria and archaea, have sparse and biased fossil records [27, 32].

Gene family histories provide a complementary source of information about the portions of the species tree that are absent from a reconstructed phylogeny. In particular, horizontal gene transfer (HGT), the exchange of genetic information across species, can connect sampled lineages to extinct or unsampled ones. Probabilistic models of gene duplication, transfer, and loss (DTL) have established that gene family histories can contain information about the structure of the species tree and the relative timing of speciation events [26, 1, 7, 23, 24]. Transfers involving ghost species were initially studied mainly as a potential source of bias: unobserved donor or recipient lineages can affect the interpretation of transfers and introgression within the sampled tree [29, 30]. More recent work has shown that HGT can instead be viewed as a useful signal to reveal the presence of ghost species along the branches of an incomplete tree [28, 25].

This signal arises because gene trees retain information about interactions that occurred outside the sampled species tree. Under a strict molecular clock, gene tree node heights provide temporal information about the events in a gene family’s history, while phylogenetic reconciliation relates these events to trajectories along the species tree. Consequently, both the topology and branch lengths of sampled gene trees (SGTs) may carry information about interactions between lineages of the sampled species tree (SST) and ghost lineages. This raises a natural theoretical question: *which features of the complete species tree (CST), containing both sampled and ghost lineages, are identifiable from sampled gene trees or from the transfers inferred from them?*

Previous work establishes that ghost lineages can affect observable gene histories and that some signatures of their presence can be detected [28, 25], but it does not characterize mathematically how much of the unobserved species tree is encoded in these signatures. Here, we address this question under idealized DTL models with an infinite number of gene families, assuming that reconstructed gene trees have the correct topology and branch lengths consistent with a molecular clock. Within this framework, and relying on simplifying assumptions summarized in Appendix A we establish four identifiability results.

1. **Ghost LTT recovery**. Transfers identify, for each branch of the sampled species tree, the ghost lineage through time (LTT) curve averaged over the time intervals delimited by the SST node heights. In particular, this determines the LTT curve of the complete tree averaged over the same subdivision.
2. **Recovery of speciation times in the complete tree**. Under exact clock dating and positive transfer rate, the atoms of the distribution of SGT node heights are exactly the internal node heights of the CST. Thus, the internal speciation times of the complete tree, including ghost speciations, are identifiable.
3. **Birth-death-sampling identifiability from hidden births**. These hidden speciation times suffice to restore identifiability of birth-death-sampling parameters, through a reduction to a recent result of Dieselhorst and Stadler [6].
4. **Generic recovery of complete species tree topology and branch lengths**. For known constant DTL rates with a positive transfer rate, the distribution of SGTs almost surely identifies the true completion and therefore uniquely determines the topology and all the branch lengths of the complete species tree.

These results show that information from gene trees can supplement reconstructed species phylogenies both when estimating speciation, extinction and sampling rates (*b, d, ρ*) in constant-rate birth-death-sampling models, and for reconstructing biodiversity through time. More generally, they provide a theoretical basis for treating the effects of ghost lineages on gene trees not only as a source of bias, but also as a source of information about otherwise unobserved diversity. These results will likely motivate the future development of dedicated phylogenomic methods for recovering ghost lineages and their evolutionary history from signals of horizontal transfer.

## 1 Model and definitions

### 1.1 The birth-death model of species evolution

Let *S* be a binary, dated, rooted species tree. We note *N* (*S*) the set of all its nodes (including leaves), and *I*(*S*) the subset of its internal nodes. For each node *u* ∈ *N* (*S*), we note *h*(*u*) for the height of node *u* (that is, the length of time that separates it from the present), and we note *h*(*S*) for the set of node heights, so that *h*(*S*) = {*h*(*u*)|*u* ∈ *N* (*S*)}. For any extant leaf *l*, we have *h*(*l*) = 0, while extinct leaves and other nodes have *h*(*l*) > 0. Let *X* be a subset of the leaves. We note *S*_|*X*_ for the reconstructed subtree of *S* whose leaves are *X*. This tree is referred to as the sampled species tree (SST).

We let *b* and *d* be two nonnegative, continuous functions of time representing the speciation and extinction rates, respectively, and let *ρ* ∈]0, 1] denote the uniform sampling probability. We assume the tree *S* is generated as a result of a constant-rate birth-death-sampling process (CRBDS) with these parameters [33].

### 1.2 The DTL model of gene evolution

We model gene evolution as a continuous-time process along *S*. This follows the standard duplication-transfer-loss (DTL) reconciliation framework, in which the discordance between a gene tree (GT) and a species tree is explained by gene duplications, horizontal transfers and losses, in addition to ordinary speciations [26, 1, 7]. We consider that all gene families originated before or at the root of the species tree. In this framework, a reconciled gene history is a GT together with a map of its nodes into the species tree and an event label for each branching or termination event.

Conditionally on the complete species tree (CST) *S*, each gene copy evolves along the species branch that contains it. On a branch alive at time *t*, and during an interval *dt*, a copy duplicates with probability *δdt* + *o*(*dt*), is lost with probability *λdt* + *o*(*dt*), and transfers with probability determined by a transfer kernel. We consider the case where each ordered pair of contemporaneous donor and receiver *species* has a transfer intensity *τ* (another transfer model, the Dated DTL model implemented in the simulator *Zombi* [5], is discussed in Appendix F). A duplication creates two copies in the same species, a loss removes the copy, and a transfer creates a descendant copy in the receiver species while the donor copy continues along the donor lineage. Gene copies are also affected by species tree events. When a gene copy reaches a speciation node of *S*, it is inherited by both descendant species. When it reaches an extinction node, the copy disappears.

A sampled gene tree (SGT) is obtained by retaining only gene copies sampled in the represented leaf set *X* and suppressing unobserved parts of the complete gene history. Internal nodes of the SGT may thus correspond to speciations in the CST (represented or hidden, see later), duplications, or transfers.

### 1.3 Ghost nodes and projections

#### Ghost nodes

When a speciation event occurs, a branch splits into two daughter lineages. In the sampled species tree (SST), only speciation events of the complete species tree (CST) with at least one sampled descendant on each daughter lineage remain visible; the others are invisible and are referred to as ghost nodes. Let *a, b* be the children of a node *n*. We distinguish three node types (Figure 1):

**Figure 1:**
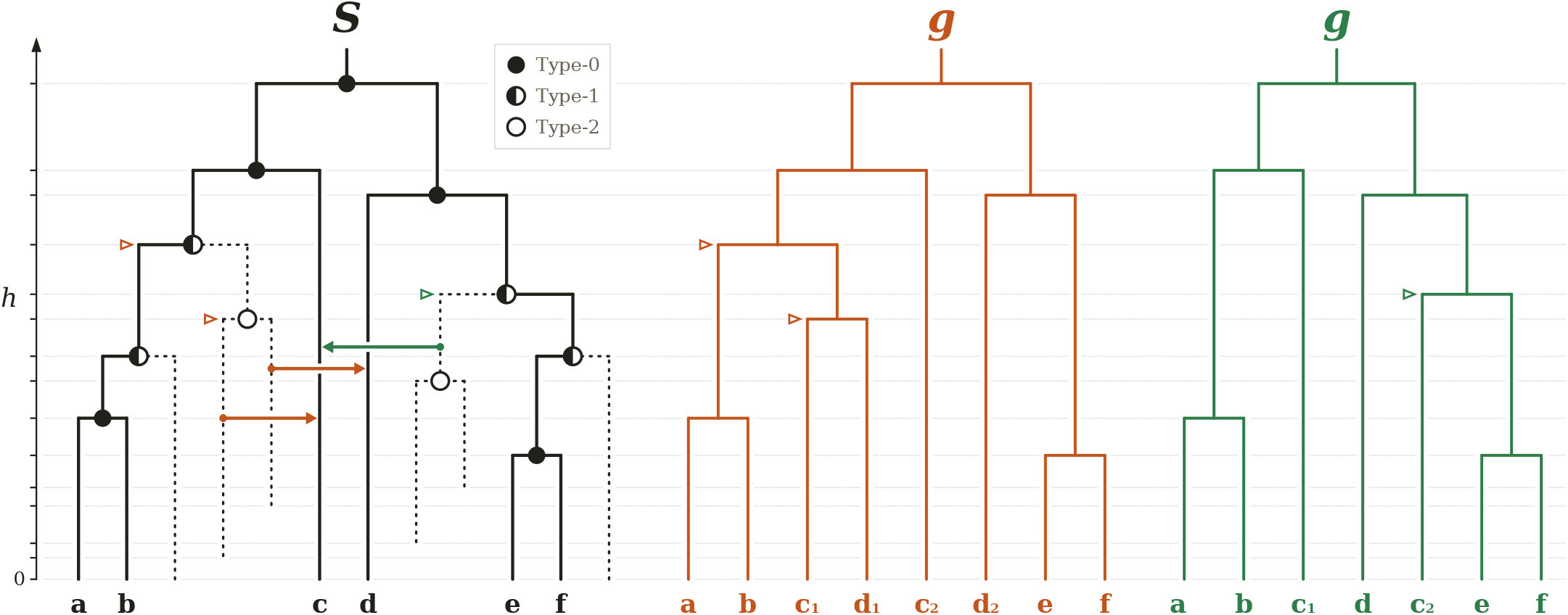
Illustration of gene-tree signals induced by transfers from ghost lineages. The tree on the left represents the complete species tree *S*, in which solid and dotted lines represent sampled and ghost branches, respectively. The three types of nodes existing in a complete species tree (see text) are depicted on the tree. Three horizontal gene transfers (colored arrows) are also indicated, each color representing a different scenario. The two transfers in orange yield the orange gene tree (center) and the transfer in green yields the green gene tree (right). The two orange transfers make visible the heights of two ghost nodes, one of type 1 and one of type 2, the latter corresponding to the MRCA of the transferred gene copies (orange triangles ▹). The green transfer makes visible a ghost node of type 1 (green triangle ▹).

1. **Sampled node (type 0)**: both *a, b* have a sampled descendant.
2. **Ghost node of type** 1: only one of *a, b* has a sampled descendant.
3. **Ghost node of type** 2: Neither of *a, b* has a sampled descendant.

#### Projections

A horizontal gene transfer that occurs in a gene family leaves a certain topological signal. For example, a transfer from a branch *a* to a branch *b*, in the absence of other events, has the effect of a subtree copy and regraft, copying the subtree on branch *b* and rebranching it on the donor branch *a*. As a result, one can deduce the donor and receiver of an HGT event from the comparison of the topology of the gene tree with that of the species tree (this is referred to as *phylogenetic reconciliation*, for a review, see [15]).

When the donor branch *a* is a ghost branch, this leads to particular signature in the gene trees [25, 29]. For instance, the donor branch being absent, the recipient branch *b* may appear regrafted at the branch closest to the ghost branch *a* in the sampled species tree. While we called this branch the *induced* branch in [25], we prefer in this article a more neutral term to help formalize the nature of the topological signal from ghost transfers on the SGTs: we simply define the *projection* of a ghost branch as the branch of the SST to which it is closest.

## 2 Recovering ghost LTTs from transfers

In a previous work [28] we showed, on both simulated data and empirical data, that ghost species, when involved as donors of horizontal gene transfers, can create an excess of apparent transfers assigned to the branch of the sampled tree nearest to the true ghost donor (the *projection* of the true donor). In this section we move from that empirical observation towards a theoretical demonstration of our ability to obtain quantitative estimates of ghost diversity through time. Under some simplifying assumptions detailed below (no loss, no duplication, and gene transfers emitted only by vertically inherited copies), we recover three complementary summaries (Figure 2): (i) branchwise ghost LTTs, giving the number of ghost lineages projecting onto each branch of the SST, averaged over each time interval defined by consecutive node heights of the SST; (ii) branchwise ghost group sizes, giving the total length of all ghost branches projecting onto each sampled-tree branch; and (iii) treewise ghost LTTs, obtained by summing the branchwise ghost LTTs across the sampled tree. Adding the known sampled-tree LTT yields the complete-tree LTT, averaged over the same intervals. This recovery requires only the observed SGTs and the SST.

**Figure 2:**
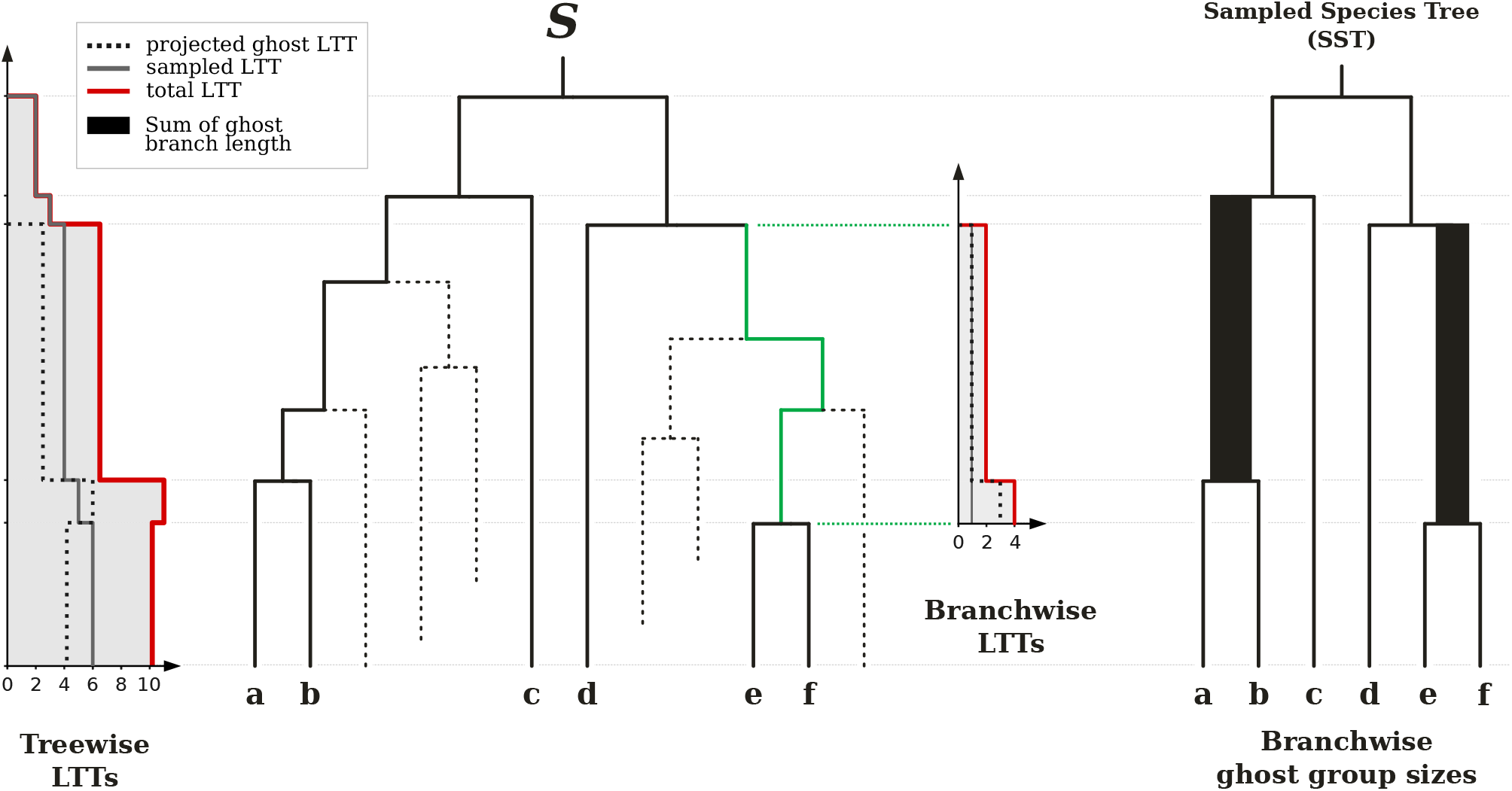
Illustration of the ghost-lineage summaries that can be recovered from sampled gene trees and the sampled species tree. *S* is the same as in Figure 1 with solid and dotted lines representing sampled and ghost branches, respectively. The treewise lineage-through-time (LTT) curves (left) give the numbers of sampled lineages (grey), ghost lineages (dotted black), and total lineages (red) alive through time, for each interval defined by nodes of the sampled species tree (nodes of type 0 in Figure 1). A branchwise LTT curve is represented for one branch of the tree, highlighted in green in *S*. For each interval defined by nodes of the sampled species tree, the number of lineages (sampled, ghost and total) projecting onto that sampled-tree branch is indicated. Right: sampled species tree. The width of each branch is proportional to the total length of the ghost branches projecting onto it, obtained by integrating its branchwise ghost LTT over time.

### 2.1 Model and assumptions

Let

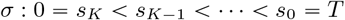

be the subdivision of time induced by the internal node heights of the SST, and write

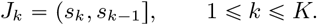

For *x* ∈(0, *T*), let *ι*(*x*) be the unique index such that *x* ∈*J*_*ι*(*x*)_. For a ghost branch *g*, let *α*_*X*_ (*g*) be the nearest ancestor of *g* with sampled descendants and set

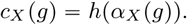

Thus *c*_*X*_ (*g*) is the height at which *g* first reconnects, backwards in time, to the part of the complete tree that survives in the SST. Its projection *π*_*X*_ (*g*) is the corresponding branch of the SST.

For an integrable function *f*, define

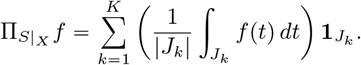

For a branch *a* of the SST, define the branchwise **projected ghost LTT**

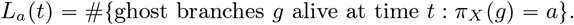

In this section, we estimate an average of this quantity on each interval of the subdivision: we hence obtain a coarse approximation of the actual *L*_*a*_ function.

For a given interval index *l*, and an SST branch *a*, we write

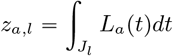

This is the total length of ghost branches descending from *a*, over the interval *J*_*l*_

#### Assumption 1

(Only vertically inherited copies can transfer). *For this section, condition on a finite CST S and its SST S*_|*X*_, *with* |*X*| ⩾ 2. *We observe N families, independent given the species tree S. Each family starts with a single copy at the root of the CST, and evolves without duplication or loss. We assume that only vertically inherited copies can be transmitted to other species, at rate τ >* 0 *(copies descending from a transferred copy are not transferred again)*.

As a result of this assumption, the number of transfers between any two species of the CST is Poisson distributed, and we will be able to more easily infer the LTTs.

Under Assumption 1, when a transfer occurs from a branch *b* of the CST to another branch *r* of the SST at time *t*, two different things can happen, depending on whether *b* is a ghost branch or is part of the *SST*. If *b* is part of the SST, then a copy of the subtree of *S*_|*X*_ below *r* is grafted onto *b*, at time *t*. If on the other hand, *b* is a ghost branch, then a copy of the subtree below *r* is grafted onto the branch *π*_*X*_ (*b*), at the height *c*_*X*_ (*b*) (independently of *t*). We can therefore identify the donor, receiver and time (*b, r, t*) for donors of the first kind, and we can identify (*π*_*X*_ (*b*), *r, c*_*X*_ (*b*)) for transfers of the second kind, and the actual value of *t* is not observed for them.

Transfer heights are continuously distributed, whereas the set of nearest ancestor heights *c*_*X*_ (*g*), for ghost branches *g*, is a finite set. Transfers coming from ghost branches are therefore aggregated on a finite set of heights, whereas transfers coming from sampled branches are continuously distributed. We can therefore separate these two kinds of transfers: “ghost” transfers are the transfers whose time is observed for several different families, whereas “nonghost” transfers almost surely happen at most once for a given height.

Let *a* be a branch of the SST, and let *r* be an interval of the subdivision. Let *Y*_*a,r*_ be the number of transfers whose donor is a ghost and projects to *a*, and whose receiver is a branch *r* of the *SST*. Let 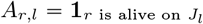. We then have:

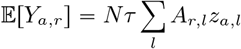

This is a linear system, the matrix *A* is known, and our goal is to invert this linear system to obtain *z*_*a,l*_ to compute ghost LTTs.

#### Proposition 1

(Projected LTT recovery). *Under Assumption 1, for every branch a of S*_|*X*_, *the distribution of SGTs identifies*,

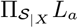

*Consequently, for every branch a of S*_|*X*_, *the total ghost branch length projecting onto a*,

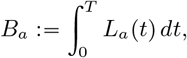

*is identifiable. Moreover, the complete-tree lineage-through-time curve, averaged over the same subdivision, is identifiable and is given by*

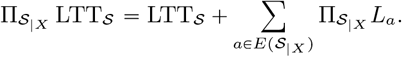

As a result, we can infer the value of the ghost LTT for every branch of the sampled phylogeny, averaged over each interval *J*_*k*_. The constant *N* is known, and *τ* can be estimated by considering the intensity of transfers between two branches of the SST: transfers between sampled branches can be separated from transfers from ghost lineages, because the corresponding MRCA height is continuously distributed, as opposed to the transfers from ghost lineages.

The proofs are given in Appendix E.

#### Branchwise LTTs

The finest object delivered by the recovery procedure is a separate ghost LTT for each branch of the sampled tree. By Proposition 1, for every *a ∈ S*_|*X*_ we identify the projected ghost LTT

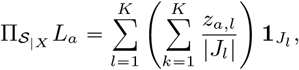

a step function on the subdivision whose value on each interval *J*_*l*_ is the time-averaged number of complete tree lineages that project onto *a* and are alive during *J*_*l*_.

#### Branchwise ghost group sizes

By integrating the ghost LTT of a branch over time, we obtain the total ghost branch length descending from this branch. For a given branch *a* of the SST, we can obtain the ghost group size *B*_*a*_ as:

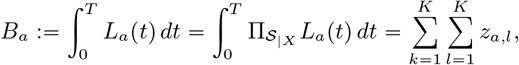

a single scalar per sampled tree branch.

#### Treewise LTTs

Summing the branchwise projected LTTs over the whole sampled tree yields the projected complete tree LTT

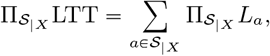

the total number of alive lineages, averaged over each interval of the subdivision. This is the global diversity-through-time curve of the complete tree, which is of interest in itself but also because it identifies the parameters of time-dependent birth-death-sampling models.

## 3 Recovering ghost nodes from finitely many gene families

Ghost nodes of types 1 and 2 can become visible in gene trees as a result of evolutionary histories involving transfers from ghost species to sampled species (Figure 1). Here we provide explicit finite-sample lower bounds on the probability of a ghost node height being visible in at least one gene family, in a simplified regime of gene evolution with no duplication and no loss. Observing a ghost node height once does not certify that it is a speciation height, however: we need to observe it in at least two distinct gene families to reach this conclusion. We give bounds for observing heights at least once in this section, and bounds for observing them at least twice in the appendix.

Assuming no loss or duplication allows us to make sure any species branch contains at least one copy at all times. Therefore, the instantaneous transfer rate between two contemporaneous species is at least *λ*, rather than 0 if some species could carry no copy. The lower bounds on the probability of observing heights count only direct transfers carried by vertically inherited copies. They are therefore conservative for the full process with transfer intensity *τ* > 0 and no loss (*λ*” 0): additional transfers create extra sampled descendants but do not erase a node that is already represented by sampled descendants on both child sides.

Let *S*_*X*_ (*t*) be the number of sampled tree branches alive at time *t*. For a ghost clade *c*, let *G*_*c*_(*t*) be the number of ghost lineages in *c* alive at time *t*, and define

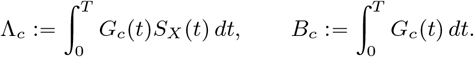

The quantity Λ_*c*_ is a measure of transfer intensity from the ghost clade *c* to the sampled tree, over the whole duration of the process of evolution. The quantity *B*_*c*_ is the total ghost length of this ghost clade, computed as the integral of the number of ghost species in *c* over the entire species history.

We illustrate the following lemma in Figure 1.

### Lemma 1

(One transfer for type 1, two transfers for type 2). *A type* 1 *ghost node can be made visible by one direct transfer from its hidden daughter clade to the sampled tree. A type* 2 *ghost node cannot be made visible by a single transfer from below that node; it is made visible when both daughter clades produce sampled descendants*.

*Proof*. Let *u* be a type 1 ghost node, with ghost daughter clade *c* and sampled daughter lineage *b*. If a vertically inherited copy in *c* transfers to a sampled tree lineage and the transferred copy leaves sampled descendants, while the copy inherited through *b* leaves sampled descendants by vertical descent, then the complete gene history contains sampled descendants from both child sides of *u*. After pruning unsampled copies, *u* has at least two sampled descendant branches and is retained as an internal node of the SGT at height *h*(*u*) (see green transfer in Figure 1).

Let *u* be a type 2 ghost node, with hidden daughter clades *c*_1_ and *c*_2_. Since neither daughter side has sampled descendants by vertical descent, observing an internal node at height *h*(*u*) requires sampled gene descendants from both *c*_1_ and *c*_2_. Suppose no transfer occurs from *c*_1_. Then all copies in *c*_1_ remain within *c*_1_, and therefore cannot leave a sampled descendant (same for *c*_2_). As a result, *u* is not realized as the MRCA of two sampled gene copies, and cannot be represented in the SGT. Conversely, if one transfer from a vertically inherited copy *c*_1_ and one transfer from a vertically inherited copy *c*_2_ both leave sampled descendants, the two sampled transferred copies have most recent common ancestor *u* in the complete gene history, and the SGT contains an internal node at height *h*(*u*) (see orange transfers in Figure 1).

### 3.1 Recovering ghost nodes of type 1

Let *c* be the ghost daughter clade of a type 1 ghost node *u*. Direct vertical-copy transfers from *c* to the sampled tree form a Poisson process with mean *τ* Λ_*c*_. Therefore, for one gene family,

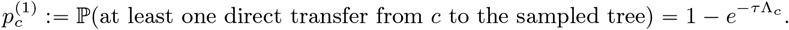

Conditional on this event the type 1 height is visible by Lemma 1, so 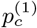 is a lower bound on the probability that the node’s height *h*(*u*) is visible in one family. If

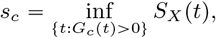

then Λ_*c*_ ≥ *s*_*c*_*B*_*c*_. For a type 1 ghost clade, *s*_*c*_ ≥ 1 because the non-ghost side has sampled descendants. Hence over *N* independent gene families,

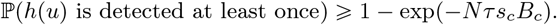

As a result, taking

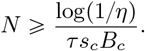

ensures detection with probability at least 1 − *η*

It remains to calibrate *τ*. We want to set this parameter in such a way that the unique vertically inherited gene copy is the donor of *κ* transfers on average. Let

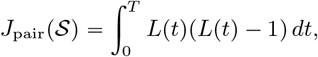

be the average number of transfer partners where *L*p*t*q is the total number of complete tree lineages alive at time *t*. If *M*_*S*_ =|*S*| is the number of complete tree branches, setting

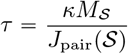

calibrates the vertical-copy process to have *κ* expected departing transfers per complete tree branch on average. With this choice of *τ*,

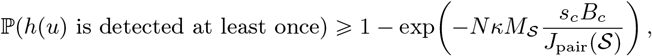

and it is enough to take

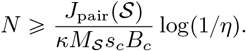

#### Example 1.

*Suppose that J*_pair_(*S*)/*M*_*S*_ = 100, *i*.*e. there are 100 times more branches in total than in the sampled tree, and calibrate to κ* =0.1 *expected departing transfers per branch. Then τ* = 10^−3^. *If s*_*c*_*B*_*c*_ =1 *(for instance, having at least* 10% *of species in the sample species tree over the period the ghost clade was alive, and having a ghost clade with sum of branch lengths above* 10*), then N* = 5000 *independent gene families give*

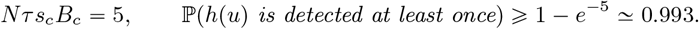

### 3.2 Recovering ghost nodes of type 2

Let *u* be a type 2 ghost node with child ghost clades *c*_1_ and *c*_2_. Let

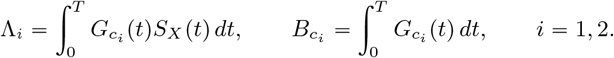

The transfer processes from *c*_1_ and *c*_2_ to the sampled tree are independent Poisson subprocesses, so that the transfer intensity between species is always *τ*. Hence the probability that both daughter sides generate at least one direct sampled transfer in one gene family is

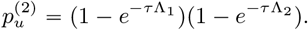

By Lemma 1, this is a lower bound on the probability that the SGT contains an internal node at height *h*(*u*). If

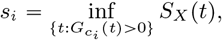

then 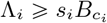. We use the following inequality, valid for all *t* ≥ 0

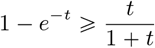

and therefore

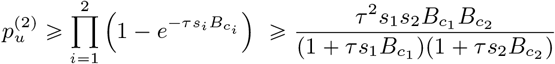

Over *N* independent gene families,

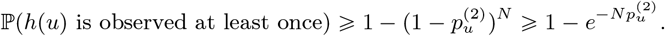

Thus 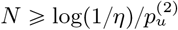 is sufficient for detection with probability at least 1 − *η*, so

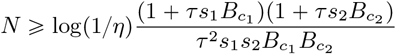

suffices.

#### Example 2.

*If, as before, we take τ* =10^−3^, *and s*_1_*B*_*c*_ = *s*_2_*B*_*c*_ = 1, *we get:*

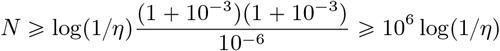

Of course, this is merely a sufficient condition to ensure we observe type 2 ghost node heights at least once with high probability, so we have not proven that we *need* that many gene families to observe these ghost node heights. On the other hand, the large lower bound is coherent with the fact that two independent, relatively rare events must happen in the same gene family for us to observe a ghost node of type 2.

## 4 Recovering internal speciation times

We demonstrated that ghost nodes of types 1 and 2 could become visible in gene trees through horizontal gene transfers. Here we go one step further and explore the possibility of recovering the node heights of the complete species tree using the distribution of node heights of sampled gene trees. Recovering all node heights of the complete species tree will be useful both to infer the parameters of the birth-death process which generated the complete species tree, and to reconstruct the latter (see section 6).

For a sampled gene tree *G*, let

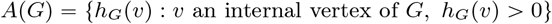

be the set of its positive internal node heights, where *h*_*G*_(*v*) denotes the height of *v*. Since the SGT is a random tree, *A*(*G*) is also a random object. We call a height *t* > 0 an *atom* of the node-height distribution if *P* (*t* ∈ *A*(*G*)) > 0.

For a branch *b* of the CST and a time *t*, let *q*_*b*_(*t*) be the probability that a copy present on branch *b* at time *t* leaves a sampled descendant.

Let *u* be a node of the CST, with child branches *b*_1_, *b*_2_. We say that the node *u* is *X*-accessible if 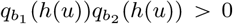. This quantity represents the probability that a copy at *b*_1_ and a copy at *b*_2_, both starting at time *h*(*u*), evolve to have sampled descendants, which would make the height *h*(*u*) visible in a sampled gene tree.

Theorem 1 below shows that the atoms are exactly the heights of the *X*-accessible nodes of the CST.

### Remark 1.

*We employ the word “atom” above, in the sense of measure theory: it is an atom of the measure*

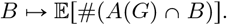

*whose atoms are exactly heights t such that P* (*t* ∈ *A*(*G*)) > 0. *To illustrate, consider a rooted tree with two leaves A, B, and root C. Then, if δ, τ, λ* ≥ 0 *and a gene originates at C, the node height C has a positive probability of being observed in the gene tree, e*.*g. if the daughter lineages on A and B do not go extinct. On the other hand, assume a copy evolves along branch A or B. The waiting time to the next event of duplication, transfer or loss has a density, so an event of duplication or transfer, which could generate a node height at some time on branch A or B has probability* 0 *to occur at any particular time t*.

### Theorem 1

(Atoms identify accessible complete-tree speciation times). *Assume bounded DTL rates, and that each family starts at the root of S. Let A*(*G*) *be the set of positive internal node heights of a sampled gene tree G*.

*Then*

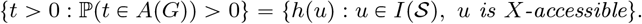

*In particular, if the internal node heights of S are pairwise distinct, the atoms of the SGT node-height distribution identify the*

*X-accessible internal nodes by their heights*.

*Proof*. Let *t* > 0 be such that ℙ (*t* ∈ *A*(*G*)) > 0.

Duplication and transfer times have a density. Therefore, among branching events, a height *t* that has positive probability of occurring in an SGT cannot have positive probability of being due to a transfer or duplication event. Therefore, *t* must be a speciation height.

We have ℙ (*t* ∈ *A*(*G*)) > 0 by hypothesis. Therefore, there exists a speciation node *u* in the CST which has a positive probability of being represented. This node is represented in a SGT if and only if both of its descendant branches *b*_1_, *b*_2_ have represented descendants. The event that both the descendant branches of a copy on *u* at time *h*(*u*) have represented descendants has probability 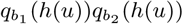, so this quantity must be positive. Hence, *u* must be *X*-accessible. Hence the inclusion

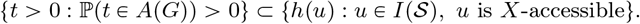

Let *u* be an internal node of the CST, and suppose that it is *X*-accessible. The probability of a copy surviving from the root of the CST to *u* at height *h*(*u*) is positive, and the probability of both copies descending from a copy on *u* at height *h*(*u*) is 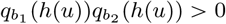. We hence have *P*(*t* ∈ *A*(*G*)) > 0.

Hence the inclusion

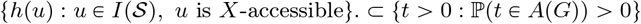

The above theorem establishes which nodes can leave a visible signature in an SGT. In particular, if the clade below *b*_1_ doesn’t transfer outside of itself, we will have *q*_*b*1_ = 0 and hence the node *u* will never appear in an SGT. Under the simplifying hypothesis of all-to-all transfers, all nodes are visible:

### Corollary 1

(All-to-all positive transfer)

*Under the pairwise all-to-all model with positive transfer between every ordered pair of contemporaneous lineages, every complete-tree internal node is X-accessible. Therefore*

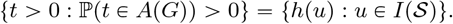

*Thus the atoms of the sampled-gene-tree node-height distribution recover all internal speciation times of the complete species tree*.

## 5 Identifiability of birth–death–sampling parameters from type 1 ghost times

Reconstructed phylogenies do not, in general, identify the birth, death and sampling parameters of the process that generated them: distinct parameter values can induce the same distribution of reconstructed trees. This non-identifiability occurs both in constant-rate models [20, 22] and in models with time-dependent rates [14]. The reconstructed tree identifies only a congruence class of models, characterized by the dLTT [14] *M*, described below, rather than the individual parameters of birth, death and sampling *b*(*t*), *d*(*t*), *ρ*.

Additional information about hidden birth events can resolve this ambiguity. Dieselhorst and Stadler [6] show that identifiability can be restored when information about these events is available along each branch of the reconstructed tree. These hidden births correspond precisely to our type 1 ghost nodes: exactly one of their two daughter lineages has sampled descendants.

In section 4, we showed that we can recover internal node heights of the complete species tree. If we are able to isolate ghost node heights of type 1 from the set of all ghost nodes heights (e.g. assuming loss rate and duplication rate are both zero, Appendix C), we can apply the results of Dieselhorst and Stadler [6] and prove identifiability of all three parameters *b*(*t*), *d*(*t*), *ρ*. Furthermore, we do not need to know the location of these ghost nodes on the sampled species tree, only their heights.

Let us recall precisely what characterizes an identifiability class of birth-death-sampling models, as in [14]. The sets of parameters which give the same distribution of reconstructed species trees can be gathered into *congruence classes*, described as follows: let *p*_0_(*t*) be the probability that a lineage alive at time *t* leaves no sampled descendants. Let us define, as in [14], the following quantities:

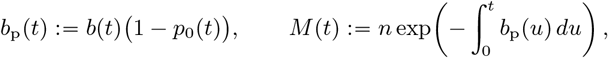

where *n* > 0 is chosen so that *M* (0) = *n* is equal to the number of sampled tips. Differentiating gives

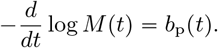

Thus, specifying *M* is equivalent to specifying *b*_p_ along with the number *n*. The quantity *M* specifies the identifiability class, but does not allow us to unambiguously identify the individual parameters *b, d, ρ*.

### Unplaced times and sampled-tree exposure

Let 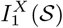 be the set of type 1 ghost nodes of the complete tree, given the sampled set of leaves *X*. Define the point measure

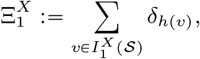

where *δ*_*t*_ is the Dirac measure at *t*.

As before, we note 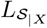 for the LTT of the SST *S*_|*X*_

#### Theorem 2

(Identifiability from type 1 times). *Consider the birth-death-sampling model θ* = (*b, d, ρ*) *of Section 1, with b and d positive and continuously differentiable and ρ* P (0, 1s, *and let T* ^*^ *be its time of origin, that is, the height of the top end of the stem of S. If two parameter values θ and* 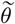 *induce the same joint distribution of* 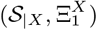, *then* 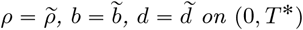.

*Proof*. Assume that *θ* = (*b, d, ρ*) and 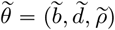 induce the same joint distribution of 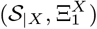, and write *β* := *b p*_*0*_, 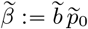. Let *n*_*X*_ (*t*) be the number of lineages of *S*_|*X*_ alive at height *t*, the stem counted, so that *n*_*X*_ = 1 on (*T*_*X*_, *T* ^*^).

Conditional on *S*_|*X*_, the hidden births on distinct branches of *S*_|*X*_, the stem included, are independent Poisson processes, each with intensity 2*β*(*t*) *dt* on the lifetime of its branch [6, Appendix A]. Summing over branches, conditional on 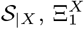 is a Poisson process on (0, *T* ^*^) with intensity

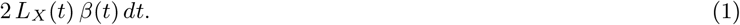

where *L*_*X*_ is the LTT of the sampled species tree. The joint distributions are equal, so the sampled-tree distributions are equal and, almost surely, the conditional distributions of 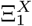 given *S*_|*X*_ are equal, hence so are the intensities (Equation (1)), almost everywhere. We have *L*_*X*_ *≥* 1 on (0, *T*^*^) and *β* is continuous, so 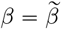 on (0, *T*^*^).

Equality of the sampled-tree distributions gives 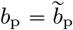 on (0, *T*^*^) [14]. Hence *b* = *b*_p_ + *β* and *p*_0_ = *β*/*b* are identified, and so is ρ = 1 − *p*_0_(0). Finally, with *t* increasing towards the past, *p*_0_ solves

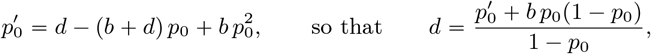

where 1 − *p*_0_ > 0 because *ρ* > 0. Hence 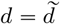 on (0, *T*^*^).

## 6 Recovering the CST topology

The previous sections demonstrate the recovery of internal node heights of the complete species tree, recovery of the LTT, or identifiability of the BDS model using the additional information provided by gene trees. However, these results provide very little information about the topology of the CST. We now prove that the distribution of SGTs almost surely identifies the CST (topology and branch lengths). Reconstructing this tree is a major goal of phylogenetics, but it is usually infeasible due to the lack of information about extinct species. The following theorem shows that in an ideal case, the distribution of SGTs provides sufficient information.

### Theorem 3

(Almost-sure identifiability of the complete topology). *Fix positive constant DTL rates θ =* (*δ, τ, λ*), *with τ* > 0. *Assume a pairwise “all-to-all” transfer model as before. Let S*^*^ *be a CST generated by* CRBDS *process, with sampled leaf set X and SST S*_|*X*_. *Suppose that the distribution of SGTs is observed. Then, almost surely, no non-isomorphic completion of S*_|*X*_ *has the same SGT distribution as S*^*^. *Equivalently, the full observed distribution of SGTs identifies the positive hidden terminal endpoint multiset, the complete internal topology and all branch lengths of S*^*^, *up to relabelling of unsampled leaves*.

*Proof*. The proof is given in Appendix D. We sketch the proof idea here.

We attribute distinct weights to the sampled leaves of the CST, and study the set of MRCA heights of pairs of leaves of an SGT. We show that the set of MRCAs that occur with positive probability are the internal nodes of the CST, and also that the density of MRCA heights generated by duplications and transfers identifies the CST LTT, which gives us the heights of extinct tips of the CST, as well as the total number of extant tips.

Finally, we show that these sets of MRCAs of two given SGT leaves allow us to reconstruct the CST topology, and that there is almost surely a single possibility, given that birth and death events on the CST are continuously distributed.

## 7 Discussion and future work

In line with previous work [28], our results support using HGT to obtain information about ghost lineages. Under ideal clock-like DTL assumptions, the signal enables the recovery of four distinct levels of hidden structure: projected ghost LTTs, internal speciation heights, birth-death-sampling parameters, and even the complete topology (with probability 1).

The number of HGT events between pairs of species permits the estimation of LTT curves at different granularities. Identifying these transfers is enabled by phylogenetic reconciliation, so that future work could estimate ghost LTT curves from already existing sequence data.

Identifying ghost nodes of type 1 seems practically feasible according to back-of-the-envelope lower bounds on the probability of observing them, which in turn would allow us to identify parameters of a general birth-death-sampling model according to Dieselhorst and Stadler [6]. Separating them from ghost node heights of type 2 is more difficult, but we think that this may not be a major obstacle: the probability of observing ghost node heights of type 2 is relatively small compared to the probability of observing ghost node heights of type 1.

Due to the mathematical complexity of the general DTL process with relaxed molecular clock, we establish results under perfect clock hypotheses, and other hypotheses when we see fit, e.g. recoverability of transfers in section 3, restriction to the case of a single transfer per family in section 2, constant DTL rates in section 6. These simplifying assumptions allow us to separate the problem of inferring events from the information that can be extracted when these events are known.

We expect that some of our conclusions will not hold in practice, for example the recoverability of all internal CST nodes or the recoverability of the CST itself. We leave the question of how to transform these theoretical results into practically usable methods for the estimation of ghost diversity for future work.

## Appendix

### A Summary of assumptions

**Table 1:** Hypotheses specific to each result.

| Result | Duplication and loss | Transfers (additive) | DTL rates |
| --- | --- | --- | --- |
| Ghost LTTs (Prop. 1) | none | all-to-all, from vertically inherited copies only | unknown |
| Visibility bounds (Sec. 3) | none | all-to-all | given |
| Speciation times (Thm. 1, Cor. 1) | allowed | all-to-all, $\tau > 0$ | unknown |
| BDS parameters (Thm. 2) | none (App. C), or allowed with known rates (Thm. 3) | all-to-all, $\tau > 0$ | unknown without duplication and loss, known otherwise |
| Complete topology (Thm. 3) | allowed | all-to-all, $\tau > 0$ | known constants |

### B Note on the molecular clock hypothesis

Our results assume that SGT node heights are measured in units of species time. If each gene family evolves with its own unknown, constant substitution rate *r*_*G*_ > 0, only the heights *h*_*G*_(*v*) = *r*_*G*_*t*_*G*_(*v*) are observed, where *t*_*G*_(*v*) is the height of the vertex *v* in units of species time. The following proposition shows that *r*_*G*_ is almost surely recovered from the SST as soon as the SGT contains two vertices coming from distinct SST speciations. We therefore keep only the families for which some pair of SGT node heights can be rescaled to coincide with a pair of SST speciation heights, and rescale them to species time; without loss and with |*X*| *≥* 3, this is every family.

#### Proposition 2

(Two-point calibration under gene-wise constant clocks)

*Let S be generated by a birth-death-sampling process with positive rates, and let H*^+^ *and* 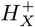 *be the sets of internal node heights of S and S*_|*X*_. *Conditionally on S, let G be an SGT generated by a DTL process with bounded rates, with substitution rate r*_*G*_ > 0. *For two internal vertices v, w of G with distinct positive observed heights, let*

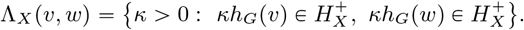

*Then, almost surely*, Λ_*X*_ (*v, w*) = {1/*r*_*G*_} *if t*_*G*_(*v*) *and t*_*G*_(*w*) *are two distinct elements of* 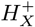, *and* Λ_*X*_ (*v, w*) = ∅ *otherwise*.

*Proof*. We first show that, almost surely, height ratios only coincide trivially: if *t*_1_, *t*_2_ are heights of internal vertices of *G* in species time and *s*_1_ ≠ *s*_2_ are elements of 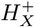, then

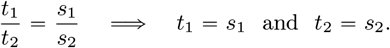

If *t*_1_, *t*_2_ ∈ *H*^+^, write *t*_1_ = *h*(*u*), *t*_2_ = *h*(*v*), *s*_1_ = *h*(*x*), *s*_2_ = *h*(*y*). Conditionally on its topology, the order of its events and its number of unsampled extant leaves, the event heights of *S* have a joint density, and unless (*u, v*) = (*x, y*) the equation *h*(*u*)*h*(*y*) = *h*(*v*)*h*(*x*) defines a null set. If *t*_1_ or *t*_2_ is the height of a duplication or a transfer, then conditionally on *S* and on the discrete part of the gene history (its skeleton), this height has a density, and the equation again has probability zero. Countable unions over topologies, skeletons and quadruples conclude.

Now let *κ* ∈ Λ_*X*_ (*v, w*), with *κh*_*G*_(*v*) = *s*_1_ and *κh*_*G*_(*w*) = *s*_2_; then *s*_1_ ≠ *s*_2_ since the observed heights are distinct. Since *h*_*G*_ = *r*_*G*_*t*_*G*_, we get *t*_*G*_(*v*)/*t*_*G*_(*w*) = *s*_1_/*s*_2_, hence *t*_*G*_(*v*) = *s*_1_, *t*_*G*_(*w*) = *s*_2_, and *κ* = *s*_1_/*h*_*G*_(*v*) = 1/*r*_*G*_. Thus Λ_*X*_ (*v, w*) is empty unless *t*_*G*_(*v*), *t*_*G*_(*w*) are distinct elements of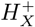, in which case it contains 1/*r*_*G*_ and nothing else.

### C Isolating ghost heights of type 1

We showed in Section 4, that it was possible to identify all node heights of the CST: when all ghost nodes are “visible” (see Section 4), they correspond to the atoms in the distribution of SGT node heights.

However, it may be useful to distinguish between ghost nodes of type 1 and 2, e.g. if we want to estimate BDS parameters as in 4. Below, we show that in some cases, it is possible to isolate ghost node heights of type 1 from the set of all ghost node heights.

#### Proposition 3

(Distinguishing type 1 and type 2 ghost heights). *Fix a finite CST S with positive branch lengths, distinct internal heights, and a nonempty sampled leaf set X. Assume that each gene family starts with one copy at the root, evolves without duplication or loss, and undergoes additive transfers with bounded positive rates between every ordered pair of contemporaneous species*.

*Let H*_*X*_ *be the set of internal SST heights and, for an SGT G, define*

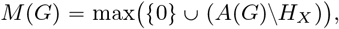

*where A*(*G*) *is its set of positive internal node heights. Then*

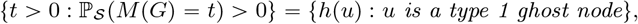

*where* ℙ_*S*_ *denotes probability conditional on S. Consequently, type 1 and type 2 ghost heights are identifiable from the SGT distribution and the SST*.

*Proof*. By Section 4, every positive atom of *M* (*G*) is a CST speciation height. Since heights in *H*_*X*_ are excluded, it must be a ghost height.

Suppose an observed gene node corresponds to a type 2 node *u*. Trace its gene ancestry from the root, and consider its first transition from a species lineage with sampled descendants to one without sampled descendants. This transition occurs either at a type 1 speciation or at a transfer. In both cases, the continuation on the represented side leaves a sampled descendant by vertical inheritance, because there is no loss and transfers are additive. The hidden continuation also leaves sampled descendants, since it leads to the observed node at *u*. Thus this earlier branching event survives in *G*. Its height is strictly greater than *h*(*u*) and lies outside *H*_*X*_ almost surely: it is either a ghost speciation height or a continuously distributed transfer time. Hence *M* (*G*) ≠ *h*(*u*), excluding all type 2 heights.

Conversely, let *u* be a ghost node of type 1. Consider the event that exactly one transfer occurs, from its hidden child branch to its represented child branch during a small interval immediately below *u*, with no other transfers anywhere in the family. Positive transfer rates and bounded total hazards give this event positive probability. Both child sides of *u* then leave sampled gene descendants, so *u* survives in *G*. The transfer node is suppressed because its donor continuation has no sampled descendant, and no other ghost node becomes visible. Therefore

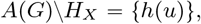

and ℙ_*S*_ (*M* (*G*) = *h*(*u*)) > 0.

This identifies all type 1 heights. The remaining ghost heights, already recovered in Section 4, are precisely those of type 2. □

### D Proofs of the main theorems

#### D.1 Proof of Corollary 1

##### Corollary 1

(All-to-all positive transfer). *Under the pairwise all-to-all model with positive transfer between every ordered pair of contemporaneous lineages, every complete-tree internal node is X-accessible. Therefore*

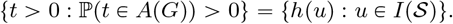

*Proof*. Let *u* be an internal node of *S* and let *c* be one of its child branches. If *c* has a sampled descendant *x*, the copy descends from *u* to *x* with no loss with positive probability, so *q*_*c*_(*h*(*u*)) > 0. If *c* has no sampled descendant, pick a height *t* inside the branch *c*. The set *X* is nonempty, so some branch *r* alive at time *t* is ancestral to a sampled leaf. The transfer rate from *c* to *r* is positive, so with positive probability the copy reaches *t*, transfers to *r* and descends to that sampled leaf, and again *q*_*c*_(*h*(*u*)) > 0. Both child branches of *u* thus have positive *q*, so *u* is *X*-accessible, and Theorem 1 gives the equality.

#### D.2 Recovering the CST topology

##### Assumptions and notation

Let *δ, λ, τ* be the duplication, loss, transfer rates respectively. Assume *τ* > 0.

For simplicity, we assume that all gene families originate at the root and the stem has length 0.

Choose distinct weights *w*_*x*_ > 0, *x* ∈ *X*, with _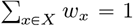_. For an SGT *G*, let *L*(*G*) be the set of its leaves, and for a copy *l*, let *σ*(*l*) be the sampled species containing *l*, and *h*_*G*_ its height function. Let

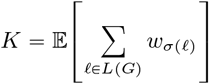

We define a measure *v* on measurable Borel sets *B* of R as:

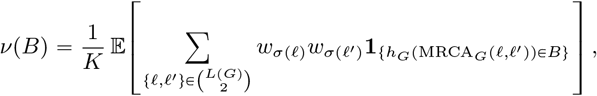

Expectations are conditional on *S*. Let *f* denote the density of the absolutely continuous part of *v*.

For *e* ∈ *E* (*t*), define

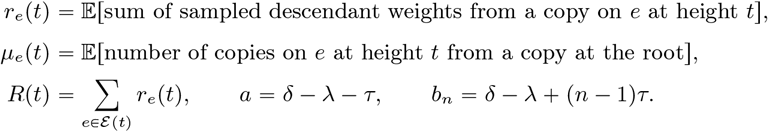

These quantities are not directly observed.

The internal speciations of *S* are denoted by *I*(*S*). We note *E*(*S*) for its set of extinction nodes. We further note *I*(*S*), *ε* (*S*) for the set of internal node heights and extinction tips of *S*. Finally, let *T* be the height of the root of *S*.

We note *U* for the open set (0, *T*)\z(*I*(*S*) ⋃ *ε* (*S*)), obtained by excluding all internal node heights and extinction heights of the SST *S*.

###### Theorem 4

(Generic identifiability of the CST). *Assume that, conditional on each finite topology and sampled set, the positive species-event heights have a joint density on each chronological chamber; the root height may instead be fixed. Then the SGT distribution determines N and, almost surely, determines S among regular completions of S*_|*X*_.

*Proof*. **1. Moment identities**. All required moments are finite: between finitely many species splittings, copy counts are dominated by a bounded-rate linear birth process. Moreover, *K* > 0 by positive root-to-sample survival.

On the open set *U*, the DTL process yields the following equations:

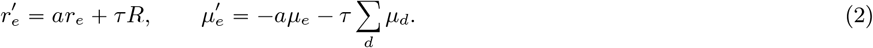

At the root node *r*, we fix *µ*_*r*_ (*T*) = 1. At leaves *l*, we fix 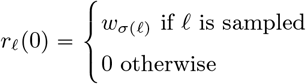

Let *c* be a branch, speciating into branches *a* and *b* at time *h* ∈ *I*(*S*). We have *µ*_*c*_(*h*) = *µ*_*a*_(*h*) = *µ*_*b*_(*h*). The differential equation defining the DTL process, as well as the boundary conditions given by the presence of a single gene copy at the root of *S* specifies the functions uniquely.

By summing the equations (2), we obtain *R*(0) = 1 and

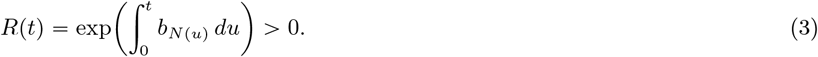

Next, we prove that at any time *t*, and for any two branches *e, f* existing at time *t*, we have *µ*_*e*_(*t*) = *µ*_*f*_ (*t*). We proceed by finite recursion over the connected components of *U* : let *e, f* be two species of an interval of *U* starting at *t*_*d*_. Notice that 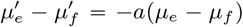 over this interval: this differential equation with boundary condition (*µ*_*e*_ − *µ*_*f*_)(*t*_*d*_) = 0 has 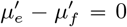 as its only solution. Speciation and extinction events all preserve the equalities between the *µ*_*i*_, so the identity is true over all of (0, *T*). We note *µ* for their common value.

Next, notice that

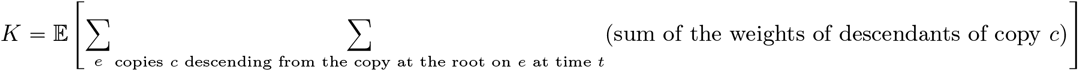

The copies at time *t* evolve independently. Therefore, by Wald’s theorem, we have

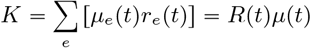

Let *h* be a speciation height, with child branches *e, d*.

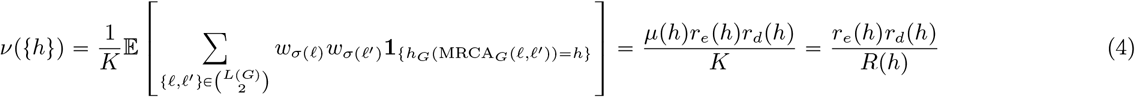

The equations of evolution for the DTL process also give the following:

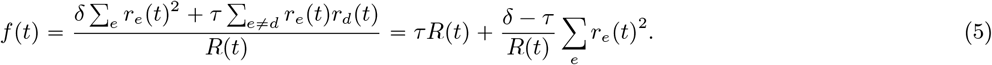

Positive branch lengths and the forcing term *τ R* >0 in (2) imply that both child *w* values at every speciation are positive. Continuous DTL event times create no other atoms, because the probability of any copy branching at a time *t* that is not a speciation time is 0.

Thus the atoms of *ν* identify exactly the speciation heights, including their maximum *T*.

##### 2. Recovery of the complete LTT

On a connected component *U* of *U*, the number *k* of contemporaneous species branches is constant. Summing 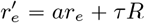 over these *k* branches gives

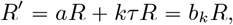

since *b*_*k*_ = *a* + *kτ*. Hence

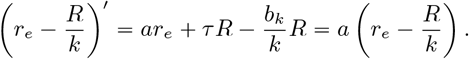

Let *t*_0_ be in this interval *U*, and let *t*_0_ be such that *u* + *t*_0_ ∈ *U*. Write

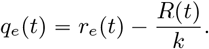

Since ∑_*e*_ *r*_*e*_p*t*q = *R*p*t*q, we have ∑_*e*_ *q*_*e*_(*t*) = 0, and hence

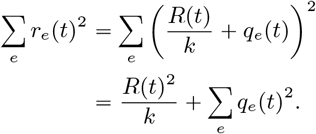

Moreover, on the considered connected component of *U*,

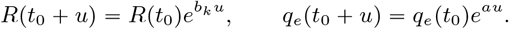

Therefore

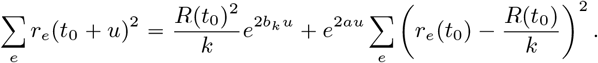

Substituting this identity into (5) gives

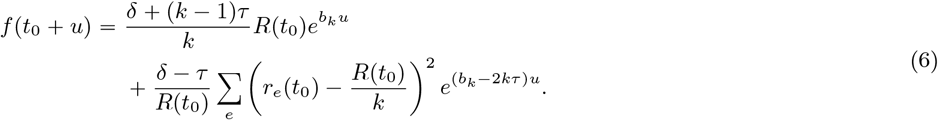

This is a sum of exponential functions with different rates. These exponential functions, restricted to any neighborhood of *t*_0_, are linearly independent (we can differentiate to obtain an invertible Vandermonde system). The larger exponential rate is *b*_*n*_, with strictly positive coefficient. We can therefore obtain the number *N* (*t*_0_) of extant species at time *t*_0_.

Moving backwards within (0, *T*), the upward jumps of *N* between successive connected components identify extinctions, and *N* (0+)− |*X*| gives the number of unsampled extant leaves. Equation (3) now determines *R*.

##### 3. Reconstruction

At height zero, initialize one component for each sampled leaf, with score *s*_*x*_ = *w*_*x*_, and *N* (0+)−p|*X*| anonymous leaf components, with score zero. Across an event-free interval of length *u*, starting at *t*, containing *n* lineages, propagate each score of currently existing branches *C* by

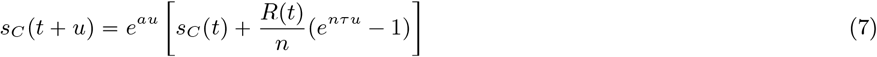

At each extinction height, add a new branch with score zero at this time. At each speciation height *h*, merge distinct branches *C, D* satisfying

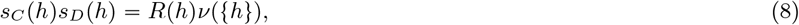

and give their parent height *h* and score *s*_*C*_ (*h*0 + *s*_*D*_ (*h*).

We proceed by induction using the terminal conditions, (2), and (4), to show that such a pair of branches (*C, D*) always exists at any speciation time.

We verify by induction over the successive species events that every current component *C* corresponding to a true branch *e* satisfies

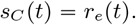

At height 0 this holds by construction: *s*_*x*_(0) = *w*_*x*_ = *r*_*x*_(0) for sampled leaves, while both scores are 0 for unsampled leaves.

If the identity holds at height *t* and (*t, t* + *u*) contains no species event, then *s*_*C*_ is also a solution of (2) with the same initial condition; hence uniqueness of solutions gives

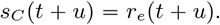

At an extinction height, the newly introduced component has score 0, which equals the *w* value of the corresponding extinct branch at its terminal endpoint.

Finally, suppose that *h* is a speciation height with true child branches *e, d*. By the induction hypothesis, the corresponding components *C*_*e*_, *C*_*d*_ satisfy

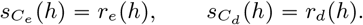

Therefore, by (4),

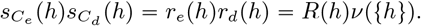

Thus the true child pair always satisfies (8). If their parent is *p*, with component *C*_*p*_, we have

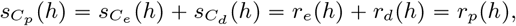

so the induction invariant is preserved.

##### 4. Almost-sure uniqueness

Fix a CST topology and a chronological ordering of its non-root species events. Let

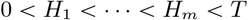

denote their heights, ordered from the present towards the root. By assumption, the random vector

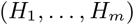

has a density with respect to Lebesgue measure. We prove that the set of event-height vectors for which the reconstruction is ambiguous has Lebesgue measure zero.

We proceed inductively through the events. Assume that all events at heights *H*_*i*_, *i* < *j* have been processed and that equal current scores occur only for equivalent components, i.e. components differing only by a permutation of anonymous leaves. Condition for the moment on fixed values

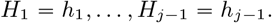

These values determine the current forest and all its scores.

Let *t*_0_ = *h*_*j*−1_ be the preceding event height, let *n* be the number of lineages on (*t*_0_, *H*_*j*_), and write *v*_*i*_ for the scores at *t*_0_. At height *t*_0_ + *u*,

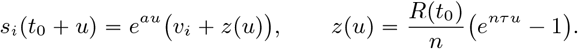

Since *τ* > 0 and *R*(*t*_0_) > 0, *z* is strictly increasing.

Suppose first that *H*_*j*_ is a speciation whose true daughter components are *i, j*. Another pair {*k, l*} gives the same product in (8) exactly when

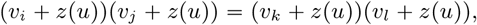

or

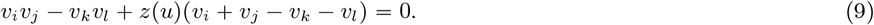

For the fixed younger heights *h*_1_, …, *h*_*j*−1_, all coefficients in (9) are fixed. If its left-hand side is not identically zero as a function of *z*, strict monotonicity of *z* shows that there is at most one value of *u*, hence at most one value of *H*_*j*_, for which this competing pair ties with the true pair.

If the left-hand side is identically zero, then

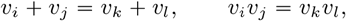

and therefore

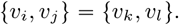

By the induction hypothesis, the two pairs are then equivalent up to permutations of anonymous components, so this does not constitute a genuine ambiguity.

After the true merger, its parent has score

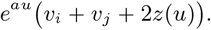

It coincides with the score of another component *k* exactly when

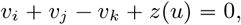

which again has at most one solution in *u*. Thus, for the fixed younger event heights, only finitely many values of *H*_*j*_ can either produce a non-equivalent competing merger or destroy the induction property on component scores.

Let *B*_*j*_ be the set of height vectors for which such an ambiguity first occurs at event *j*. For every fixed choice of the younger heights outside the previously excluded sets, the section of *B*_*j*_ in the *H*_*j*_ coordinate is finite, hence has one-dimensional Lebesgue measure zero. The older heights *H*_*j*+1_, …, *H*_*m*_ are irrelevant to this calculation. Fubini’s theorem therefore gives

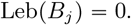

Induction over the finitely many non-root events yields

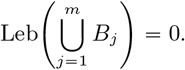

Since (*H*_1_, …, *H*_*m*_) has a joint density on the chronological chamber,

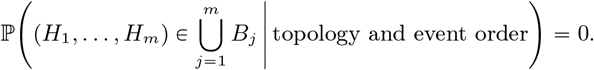

At the root only two components remain, so the final merger is forced. Hence, conditional on the fixed topology and event order, the reconstruction is almost surely unique up to relabelling anonymous leaves. Finally, the collection of finite topologies and chronological orders is countable, so the same conclusion holds unconditionally.

##### 5. Conclusion

The reconstruction of Step 3 only uses quantities determined by the SGT distribution and the known rates: the atoms of *ν*, the lineage count *n* (Step 2), and *R* (equation (3)). Let *S*’ be a completion of *S*_|*X*_ with the same SGT distribution as *S*, and assume that *S* lies outside the null set of Step 4. Then *S* and *S*’ have the same lineage count, hence the same extinction heights and unsampled extant leaves. By induction over the events, assume that the reconstructed forest coincides with both the forests of *S* and *S*’ below some speciation height *h*. The scores then equal the *r* values of both trees (Step 3), so the true child pairs of both trees satisfy (8) at *h*. By Step 4, this pair is unique up to equivalent components, so both trees perform the same merger at *h*. Hence *S*’ = *S* up to relabelling of unsampled leaves.

### E Derivation of projected LTT recovery

As before, we write *S* for the complete tree, and 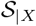 for the reconstructed tree on the leaves *X*. The present is at time 0, and the root node of the tree is at time *T* > 0.

Let

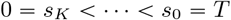

be the set of node heights in the sampled tree. For all *k* ∈ [1, *K]*, define

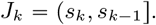

These intervals generate a finite *σ*-algebra, which we denote by 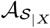. By analogy with conditional expectations, for an integrable function *f* we write

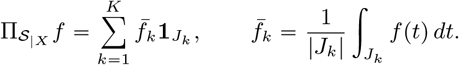

For a sampled tree branch *a* and an interval *J*_*k*_, let *L*_*a*_(*t*) be the number of complete tree lineages that are alive at time *t*. Define

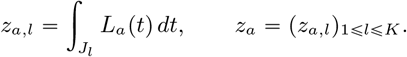

For simplicity, assume that the transfer intensity between any pair of contemporaneous species is *τ*. Let *R* be the set of sampled tree branches that can be receivers. For *r* ∈ *R*, define

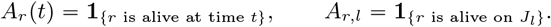

The function *A*_*r*_ is 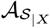-measurable. Let *Y*_*a,r*_ be the number of observed transfers, across *N* independent gene families, with induced donor *a*, receiver *r*, and induced-donor coalescence interval *J*_*k*_. Conditional on the complete tree,

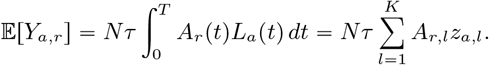

Equivalently, the count has Poisson distribution

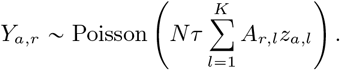

Let

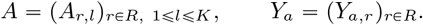

Then

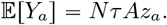

#### Lemma 2.

*The matrix* **A** *has full column rank*.

*Proof*. Let 2 ⩽*k* ⩽*K*. Choose a branch of *S*_|*X*_ with top node at height *s*_*k*−1_. It is alive during *J*_*k*_ and not on any interval *J*_*i*_ with *i* < *k*, so it corresponds to a row of *A* with zeros for indices 1 ⩽*i* < *k*, and a one for index *k*. On *J*_1_, there is only the stem, so we obtain a row where the first coefficient is 1 and the others are 0.

Such a row exists for all *k*, so by keeping only one such row for each value of *k*, we obtain a submatrix of *A* that is upper diagonal with ones on the diagonal. This matrix has rank *K*, so *A* also has rank *K*.

Since *A* has full column rank, this linear system can be inverted:

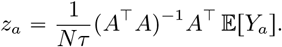

The projected LTT along induced donor *a* is therefore

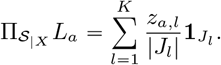

Finally, summing over sampled tree branches gives the projected complete tree LTT:

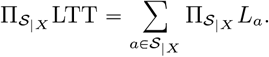

We close with the finite-sample behaviour of this inversion.

#### Proposition 4

(Unbiased estimator, covariance and concentration). *Set*

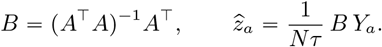

*Then*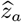 *is unbiased*, 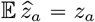, *with covariance*

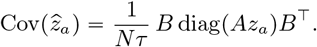

*In particular, by Chebyshev’s inequality*,

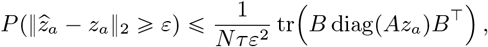

*and, writing b*_*j*_ *for the jth row of B, the jth coordinate satisfies the Bernstein bound*

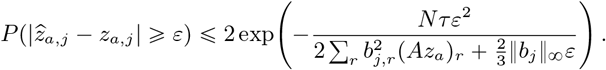

*Proof*. By linearity, 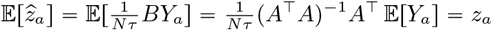, so the estimator 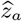 is unbiased.

Conditional on the complete tree, the random variables *Y*_*a,r*_ are independent Poisson variables, so Cov(*Y*_*a*)_ =*Nτ* diag(*Az*_*a*_).

We have

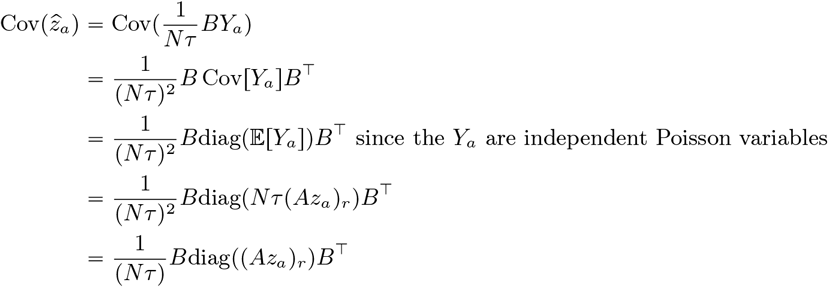

Fix *j* ∈{1, …, *K*}. For every *r*∈*ℛ*, set *m*_*r*_: E*Y*_*a,r*_*=Nτ(Az*_*a)r*_. Each *m*_*r*_ is the mean of a Poisson variable, so *m*_*r*_⩽0. The *j*th coordinate of 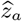 equals 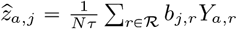. The estimator is unbiased, so 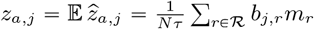.Set

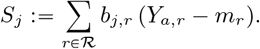

Subtracting the two previous expressions gives 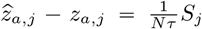.Set 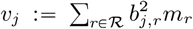 and *k*_*j*_ :=║*b*_*j*_║∞/3. Every *m*_*r*_ is nonnegative, so *v*_*j*_ ⩾ 0. The *j*th row of *BA* equals *b*_*j*_ *A*, and *BA*” *I*, so *b*_*j*_ *A* is the *j*th row of *I*. This row is nonzero, so *b*_*j*_ is nonzero. A nonzero vector has positive supremum norm, so ║*b*_*j*_║ ∞>0. Hence *κ*_*j*_ > 0.

For every *m* ⩾ 0, every Poisson variable *Y* with mean *m* and every *µ* ∈ ℝ satisfy E*e* ^µ(*Y*−*m*)^=*m*(*e*^µ^ −1−^µ^). Fix *λ* ∈ ℝ. The variables *Y*_*a,r*_, *r* ∈ *ℛ*, are independent, so the variables 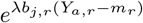, *r* ∈ *ℛ*, are independent, and the expectation of their product equals the product of their expectations. For every *r* ∈ *ℛ* apply the Poisson identity with *µ* = *λb*_*j,r*_, *m* =*m*_*r*_ and *Y* =*Y*_*a,r*_ :

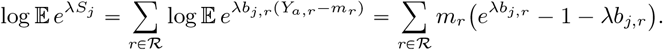

Let *u* ∈ ℝwith |*u*| < 3. For every integer *p ≥*2, the product *p*! 2.3… *p* contains the *p* − 2 factors 3, …, *p*, each at least 3, so *p*! ≥ 2.3^p−2^. The ratio |*u*|/3 is smaller than 1, so the geometric series ≥ _*q≥*0_(u|/3)^*q*^ converges to 1/1 −|*u*|/3). The exponential series, the bound *u*^*p*^ *≤*|*u*|^*p*^ and the factorial bound give

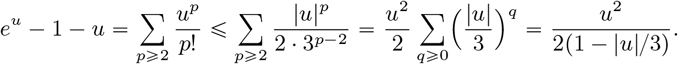

Assume now that 0 < *λ* < 1/*κ*_*j*_. Fix *r* ∈ ℛ. Then |*λb*_*j,r*_ | *≤λ* ∥*b*_*j* ∥_ = 3*λκ*_*j*_ < 3. Apply the elementary inequality with *u* =*λb*_*j,r*_ :

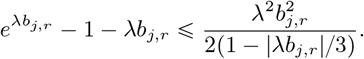

The definition of *κ*_*j*_ gives |*b*_*j,r*_ |/3 *≤κ*_*j*_, so 1 − |*λb*_*j,r*_ |/3 ≥ 1 −*κ*_*j*_*λ* > 0. Hence 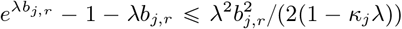. The last inequality holds for every *r* ∈ ℛ. Multiply it by *m*_*r*_ *≤* 0, sum over ∈ ℛand insert the sum into the expression of 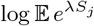

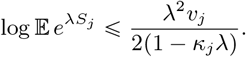

The variable −*S*_*j*_ equals ∑_*r*∈*ℛ*_ (−*b*_*j,r*_)(*Y*_*a,r*_ − *m*_*r*_). The row −*b*_*j*_ has the same supremum norm and the same squared entries as *b*_*j*_, so the same argument gives 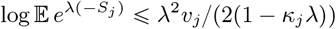. Both bounds hold for every 0 < *λ* < 1/*κ*_*j*_.

Let *v* ⩾ 0 and *c* > 0, and let *S* be a random variable such that, for every 0 < *λ* < 1/*c*, log E *e*^*λS*^ ⩽ *λ*^2^*v*/(2(1 − *cλ*)). Section 2.4 of [2] then gives, for every *t* > 0,

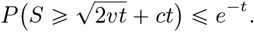

Fix *x* > 0. The constants satisfy *v* + *cx* > 0. Set *t* := *x*^2^/(2(*v* + *cx*)) and *s* := *v*/(*v* + *cx*). Then *t* > 0 and *s* ∈ [0, 1]. Direct computation gives 2*vt = sx*^2^ and *ct = x (*1 – *s) /*2, so 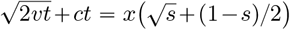. The inequality 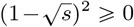 gives 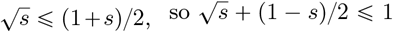. Hence 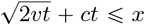, and the event {*S* ⩾ *x*} is contained in the event 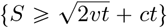. The cited bound gives

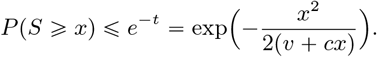

The variables *S*_*j*_ and −*S*_*j*_ satisfy the hypothesis of this bound with *v* = *v*_*j*_ and *c = κ*_*j*_. For every *x* > 0, the event {|*S*_*j*_ | ⩾ *x*} is the union of {*S*_*j*_ ⩾ *x*} and {−*S*_*j*_ *⩾ x*}, so the union bound and the bound above give

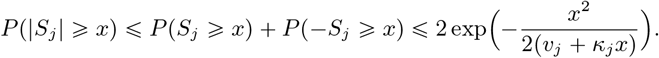

Take *x* := *Nτε*. The identity 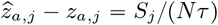 gives 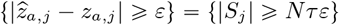. The definitions of *v*_*j*_, *κ*_*j*_ and *m*_*r*_ give

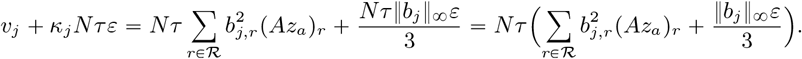

Hence

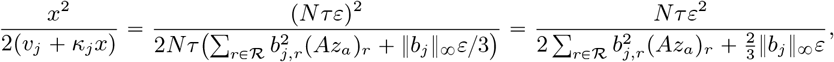

and the two-sided bound becomes

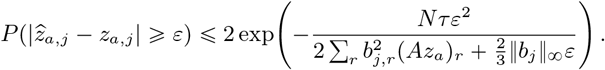

### F Recovering ghost nodes of type 1: the normalized transfer probability case

This appendix records an alternative version of the type 1 visibility calculation given in Section 3. Here the transfer intensity is *normalized* by the number of contemporaneous receivers *S*(*t*) +*G*(*t*) −1, so that each donor lineage transfers at total rate *τ* shared among the possible receivers.

The main text (Section 3) works with the pairwise model, where *τ* is calibrated so that the expected number of transfers over the whole complete tree is realistic; the normalized model recorded here is included for comparison and yields the same qualitative conclusion.

A ghost of type 1 can appear in an SGT as the result of a single gene transfer event, from the ghost to the sampled tree. We give below a lower bound on the probability of a transfer occurring from a ghost clade to the sampled tree.

For all *t* ∈[0,*T*], we let *S*(*t*) and *G*(*t*) be the number of sampled and ghost species at time *t* respectively.

We let 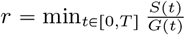 and 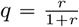.

Let 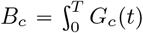 *dt* be the total branch length of ghost group *c*. We assume that we observe transfers over *N* gene families.

The intensity of transfers from group *c* to the sampled tree is

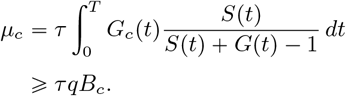

The probability of no transfer occurring from the ghost group *c* to the sampled tree is therefore smaller than exp(−*N τqB*_*c*_q).

#### Remark 2.

*This bound counts only transfers emitted by vertically inherited copies, which are present in every species at all times in the absence of duplication and loss*.

#### Example 3.

*Let c be a ghost group. Let us take the values*

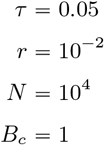

*Then we have* 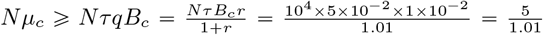, *and hence, the probability of c being observed is greater than* 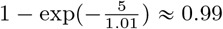.

This back-of-the-envelope calculation demonstrates that there is a regime in which ghost nodes of type 1 manifest themselves with high probability.

### G Seeing a ghost node twice

Section 3 bounds the probability that a ghost node height is observed in at least one gene family. In a general DTL history, a height produced by a duplication or a transfer is almost surely not repeated across families (Theorem 1), so a height observed in two families is almost surely a speciation height of the CST, which is what certifies it. If the height is visible in each family independently with probability at least *p*, the number of families in which it is visible dominates a binomial variable with parameters (*N, p*), whose probability of being at most 1 is

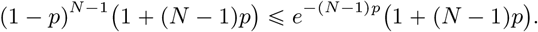

Hence

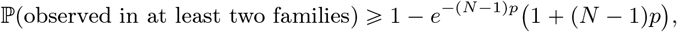

and since (1 + *x*)*e*^*−x*^ ⩽ 2*e*^*−x/2*^ for *x* ⩾ 0, it suffices that

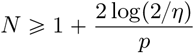

to reach probability 1 − *η*: about twice the number of families needed to see the height once. For a type 1 node one takes 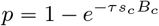, and for a type 2 node 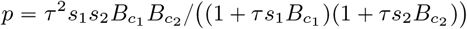, the per-family lower bounds of Section 3. In the type 1 example of that section (*τs*_*c*_*B*_*c*_ = 10^−3^, *N* = 5000), the height is observed at least twice with probability about 0.96.

## Glossary

Complete species tree.: The complete species tree is the dated species tree generated by the birth-death process.
Sampled species tree.: The sampled species tree is the reconstructed subtree induced by the sampled extant leaves *X*.
Ghost species.: A ghost species is a species lineage that is absent from the sampled species tree, either because it is extinct without sampled descendants or because it is extant but unsampled.
Ghost node of type 1.: A ghost node of type 1 is a speciation node of the complete species tree for which exactly one daughter lineage has a sampled descendant.
Ghost node of type 2.: A ghost node of type 2 is a speciation node of the complete species tree for which neither daughter lineage has a sampled descendant.
Projection *π*_*X*_.: Let *b* be a branch of the complete species tree *S*. Let *X* be a nonempty subset of the leaves of *S*. The projection *π*_*X*_ (*b*) of branch *b* onto the species tree *S* _|*X*_ is, informally, the branch of *S* _|*X*_ that contains the last common ancestor between the sampled species X and the branch b.
Completion.: A completion of a sampled tree *S*_|*X*_is a dated, rooted binary tree ℛ admitting a leaf subset *X* with ℛ _| *x*_ ≃ *S*_| *x*_, that is, a complete tree containing the observed sampled tree.
LTT: The LTT (Lineage Through Time) plot of a tree is the function that gives the number of alive species at a time *t* [19, 10].
Horizontal gene transfer.: A horizontal gene transfer is the transfer of a gene copy from one species to another contemporaneous species. It creates a new copy in the genome of the receiving species. In this paper transfers are *additive* : the new copy is added to the receiving genome. A transfer is said to be a *replacement* transfer if the received copy replaces a copy already present in the receiving genome, which is equivalent to an additive transfer followed by a loss in the receiving genome.

## Abbreviations

We use the following abbreviations throughout.

SST, CST: sampled, complete species tree
GT: gene tree
SGT, CGT: sampled, complete gene
RGT: reconciled gene tree
CRGT: complete reconciled gene tree
CRBD: constant-rate birth-death
CRBDS: consssastant-rate birth-death-sampling

## References

[1] Mukul S. Bansal, Eric J. Alm, and Manolis Kellis. Efficient algorithms for the reconciliation problem with gene duplication, horizontal transfer and loss. Bioinformatics, 28(12):i283–i291, 2012.

[2] Stéphane Boucheron, Gábor Lugosi, and Pascal Massart. Concentration Inequalities: A Nonasymptotic Theory of Independence. Oxford University Press, 2013.

[3] Arthur D. Chapman. Numbers of living species in australia and the world. Technical report, Australian Biological Resources Study, Canberra, Australia, 2009. 2nd edition.

[4] Thomas P. Curtis, William T. Sloan, and Jack W. Scannell. Estimating prokaryotic diversity and its limits. Proceedings of the National Academy of Sciences of the United States of America, 99(16):10494–10499, 2002.

[5] Adrián A. Davín, Théo Tricou, Eric Tannier, Damien M. de Vienne, and Gergely J. Szöllősi. Zombi: a phylogenetic simulator of trees, genomes and sequences that accounts for dead lineages. Bioinformatics, 36(4):1286–1288, 2020.

[6] Tobias Dieselhorst and Tanja Stadler. Information on hidden birth events restores identifiability in phylodynamic inference. https://arxiv.org/abs/2604.17926, 2026. arXiv:2604.17926 [q-bio.PE].

[7] Jean-Philippe Doyon, Vincent Ranwez, Vincent Daubin, and Vincent Berry. Models, algorithms and programs for phylogeny reconciliation. Briefings in Bioinformatics, 12(5):392–400, 2011.

[8] Terry L. Erwin. Tropical forests: Their richness in coleoptera and other arthropod species. The Coleopterists Bulletin, 36(1):74–75, 1982.

[9] Tracy A. Heath, John P. Huelsenbeck, and Tanja Stadler. The fossilized birth-death process for coherent calibration of divergence-time estimates. Proceedings of the National Academy of Sciences of the United States of America, 111(29):E2957– E2966, 2014.

[10] Andrew J. Helmstetter, Sylvain Glémin Jos Käfer, Rosana Zenil-Ferguson, Hervé Sauquet Hugo de Boer, Léo-Paul M. J. Dagallier, Nathan Mazet, Eliette L. Reboud, Thomas L. P. Couvreur, and Fabien L. Condamine. Pulled diversification rates, lineages-through-time plots, and modern macroevolutionary modeling. Systematic Biology, 71(3):758–773, 2022.

[11] Brendan B. Larsen, Elizabeth C. Miller, Matthew K. Rhodes, and John J. Wiens. Inordinate fondness multiplied and redistributed: the number of species on earth and the new pie of life. The Quarterly Review of Biology, 92(3):229–265, 2017.

[12] Kenneth J. Locey and Jay T. Lennon. Scaling laws predict global microbial diversity. Proceedings of the National Academy of Sciences of the United States of America, 113(21):5970–5975, 2016.

[13] Stilianos Louca, Florent Mazel, Michael Doebeli, and Laura Wegener Parfrey. A census-based estimate of earth’s bacterial and archaeal diversity. PLOS Biology, 17(2):e3000106, 2019.

[14] Stilianos Louca and Matthew W. Pennell. Extant timetrees are consistent with a myriad of diversification histories. Nature, 580:502–505, 2020.

[15] Hugo Menet, Vincent Daubin, and Eric Tannier. Phylogenetic reconciliation. PLOS Computational Biology, 18(11):1–29, 11 2022.

[16] Camilo Mora, Derek P. Tittensor, Sina Adl, Alastair G. B. Simpson, and Boris Worm. How many species are there on earth and in the ocean? PLOS Biology, 9(8):e1001127, 2011.

[17] Hélène Morlon, Stéphane Robin, and Florian Hartig. Studying speciation and extinction dynamics from phylogenies: addressing identifiability issues. Trends in Ecology & Evolution, 37(6):497–506, 2022.

[18] David M. Raup. Extinction: Bad Genes or Bad Luck? W. W. Norton & Company, New York, 1991.

[19] Tanja Stadler. Lineages-through-time plots of neutral models for speciation. Mathematical Biosciences, 216(2):163–171, 2008.

[20] Tanja Stadler. On incomplete sampling under birth-death models and connections to the sampling-based coalescent. Journal of Theoretical Biology, 261(1):58–66, 2009.

[21] Tanja Stadler. Sampling-through-time in birth-death trees. Journal of Theoretical Biology, 267(3):396–404, 2010.

[22] Tanja Stadler and Mike Steel. Swapping birth and death: Symmetries and transformations in phylodynamic models. Systematic Biology, 68(5):852–858, 2019.

[23] Gergely J. Szöllősi, Bastien Boussau, Sophie S. Abby, Eric Tannier, and Vincent Daubin. Phylogenetic modeling of lateral gene transfer reconstructs the pattern and relative timing of speciations. Proceedings of the National Academy of Sciences of the United States of America, 109(43):17513–17518, 2012.

[24] Gergely J. Szöllősi, Wojciech Rosikiewicz, Bastien Boussau, Eric Tannier, and Vincent Daubin. Efficient exploration of the space of reconciled gene trees. Systematic Biology, 62(6):901–912, 2013.

[25] Eric Tannier, Théo Tricou, Syrine Benali, and Damien M. de Vienne. HGTs are not SPRs: in the presence of ghost lineages, series of horizontal gene transfers do not result in series of subtree pruning and regrafting. Molecular Biology and Evolution, 42(6):msaf128, 2025.

[26] Ali Tofigh, Michael Hallett, and Jens Lagergren. Simultaneous identification of duplications and lateral gene transfers. IEEE/ACM Transactions on Computational Biology and Bioinformatics, 8(2):517–535, 2011.

[27] Alexandru M. F. Tomescu, Ashley A. Klymiuk, Kelly K. S. Matsunaga, Alexander C. Bippus, and Glenn W. K. Shelton. Microbes and the fossil record: Selected topics in paleomicrobiology. In Christon J. Hurst, editor, Their World: A Diversity of Microbial Environments, pages 69–169. Springer International Publishing, 2016.

[28] Théo Tricou, Enzo Marsot, Bastien Boussau, Eric Tannier, and Damien M. de Vienne. Gene flow can reveal ghost lineages. Evolutionary Journal of the Linnean Society, 4(1):kzaf014, 2025.

[29] Théo Tricou, Eric Tannier, and Damien M. de Vienne. Ghost lineages can invalidate or even reverse findings regarding gene flow. PLOS Biology, 20(9):e3001776, 2022.

[30] Théo Tricou, Eric Tannier, and Damien M. de Vienne. Ghost lineages highly influence the interpretation of introgression tests. Systematic Biology, 71(5):1147–1158, 2022.

[31] Kate Truman, Timothy G. Vaughan, Alex Gavryushkin, and Alexandra Sasha Gavryushkina. The fossilized birth-death model is identifiable. Systematic Biology, 74(1):112–123, 2025.

[32] Frances Westall and Keyron Hickman-Lewis. Fossilization of bacteria and implications for the search for early life on earth and astrobiology missions to mars. In Vera M. Kolb, editor, Handbook of Astrobiology. CRC Press, 2018.

[33] Z Yang and B Rannala. Bayesian phylogenetic inference using dna sequences: a markov chain monte carlo method. Molecular Biology and Evolution, 14(7):717–724, 07 1997.

